# Human metapneumovirus P protein independently drives phase separation and recruits N protein to liquid-like inclusion bodies

**DOI:** 10.1101/2021.09.24.461765

**Authors:** Kerri Beth Boggs, Nicolas Cifuentes-Munoz, Kearstin Edmonds, Farah El Najjar, Conny Ossandón, McKenna Roe, Carole L. Moncman, Trevor Creamer, Rebecca Ellis Dutch

## Abstract

Human metapneumovirus (HMPV) inclusion bodies (IBs) are dynamic structures required for efficient viral replication and transcription. The minimum components needed to form IB-like structures in cells are the nucleoprotein (N) and the tetrameric phosphoprotein (P). HMPV P binds to two versions of N protein in infected cells: C-terminal P residues interact with oligomeric, RNA-bound N (N-RNA), and N-terminal P residues interact with monomeric N (N^0^) to maintain a pool of protein to encapsidate new RNA. Recent work on other negative-strand viruses has suggested that IBs are liquid-like organelles formed via liquid-liquid phase separation (LLPS). Here, HMPV IBs in infected or transfected cells were shown to possess liquid organelle properties, such as fusion and fission. Recombinant versions of HMPV N and P proteins were purified to analyze the interactions required to drive LLPS *in vitro*. Purified HMPV P was shown to form liquid droplets in the absence of other protein binding partners, a novel finding compared to other viral systems. Removal of nucleic acid from purified P altered phase separation dynamics, suggesting that nucleic acid interactions also play a role in IB formation. HMPV P also recruits monomeric N (N^0^-P) and N-RNA to IBs *in vitro*. These findings suggest that, in contrast to what has been reported for other viral systems, HMPV P acts as a scaffold protein to mediate multivalent interactions with monomeric and oligomeric HMPV N to promote phase separation of IBs.

**IMPORTANCE:** Human metapneumovirus (HMPV) is a leading cause of respiratory disease among children, immunocompromised individuals, and the elderly. Currently, no vaccines or antivirals are available for treatment of HMPV infections. Cytoplasmic inclusion bodies (IBs), where HMPV replication and transcription occur, represent a promising target for the development of novel antivirals. The HMPV nucleoprotein (N) and phosphoprotein (P) are the minimal components needed for IB formation in eukaryotic cells. However, interactions that regulate the formation of these dynamic structures are poorly understood. Here, we showed that HMPV IBs possess the properties of liquid organelles and that purified HMPV P phase separates independently *in vitro*. Our work suggests that HMPV P phase separation dynamics are altered by nucleic acid. We provide strong evidence that, unlike results reported from other viral systems, HMPV P alone serves as a scaffold for multivalent interactions with monomeric (N^0^) and oligomeric (N-RNA) HMPV N for IB formation.

## INTRODUCTION

Human metapneumovirus (HMPV), discovered in 2001, is a leading cause of severe respiratory tract infections in infants, the elderly, and immunocompromised individuals (1). Five to twenty percent of hospitalizations from respiratory infections in young children are due to HMPV (2, 3). Symptoms of HMPV infection are similar to respiratory syncytial virus (RSV) and include fever, cough, rhinorrhea, croup, bronchiolitis, pneumonia, and asthma exacerbation (4). HMPV and RSV are members of the *Pneumoviridae* family, a viral family which was created in 2016 and classified within the *Mononegavirales* order (5). Currently, no vaccines or antiviral treatments are approved to treat HMPV infections, so most patients are managed with supportive care (4). The recent discovery of HMPV highlights the need to understand the basic mechanisms of its life cycle. Specifically, analyzing the process of HMPV replication may be crucial for identifying new targets for antiviral development.

Along with the pneumoviruses HMPV and RSV, other relevant human pathogens within the *Mononegavirales* order include Ebola virus, measles virus (MeV), and rabies virus (RABV), which have negative-sense, single-stranded RNA genomes. Though these viruses are classified within different families, they have all been reported to form membrane-less cytoplasmic structures within infected cells known as inclusion bodies (IBs) (6–9). For some negative-sense, single-stranded RNA viruses, including HMPV, IBs have been shown to house active viral replication and transcription (10–18). These processes involve several viral proteins, such as the large RNA-dependent RNA-polymerase (L), phosphoprotein (P), and nucleoprotein (N). Further analysis of these structures has shown that RSV, MeV, RABV, and vesicular stomatitis virus (VSV) IBs possess the properties of liquid organelles formed via liquid-liquid phase separation (LLPS) (11, 19–22). LLPS is a physical process by which a homogenous fluid separates into two distinct liquid phases (23). Phase separation plays a role in the formation of a variety of membrane-less cellular compartments, such as processing bodies (P-bodies), stress granules, and nucleoli, to concentrate specific proteins and nucleic acids, particularly RNA (24). Properties that define these structures as liquid organelles include the ability to undergo fusion and fission, rapid diffusion of internal contents, and a spherical shape due to surface tension (25). Though LLPS has been shown to play a role in the formation of IBs for some viruses, the physical mechanisms and materials that mediate this process in the viral life cycle are still poorly understood.

For RSV, HMPV, MeV, and RABV, the minimum viral components required to reconstitute IB-like structures in cells are the N protein, which encapsidates the RNA genome, and the P protein, which acts as a cofactor to mediate interactions between N and L (11, 20, 26, 27). VSV also requires the presence of the L protein with the N and P proteins to form IBs (19).These findings suggest that interactions between the N and P proteins regulate phase separation to form IBs as a structural platform for viral replication and transcription. Most studies thus far have focused on cellular experiments to investigate viral IB liquid dynamics, but recent publications on MeV and RSV have shown the importance of utilizing purified protein systems to analyze interactions between the N and P proteins *in vitro* (21, 22). For MeV, the purified P protein and monomeric N protein failed to phase separate independently but formed liquid droplets when mixed, similar to the requirements for IB formation observed in cells. Interestingly, when RNA was added to MeV N/P liquid droplets, it incorporated into the droplets and led to the formation of nucleocapsid-like particles that were detected by electron microscopy (21). *In vitro* experiments with RSV proteins showed that RNA-bound N protein rings and P protein form phase separated liquid droplets when combined in solution (22). These findings support the model that viral IBs form via LLPS, and this mechanism is highly dependent upon interactions between the N and P proteins. This process may enhance viral replication and transcription for RSV.

The HMPV life cycle begins with the virus attaching and fusing to a target cell to release its ribonucleoprotein into the cytoplasm. The ribonucleoprotein structure protects the genome from host nucleases and acts as a template for the L protein. The genome is used to generate capped and poly-adenylated viral mRNA transcripts that are translated by the host cell ribosomal machinery. The genome is also replicated to make positive-sense antigenome copies that can then be used to generate more negative-sense genome to package into new virions. The P protein acts as an adaptor to regulate interactions between the polymerase and RNA template during transcription and replication. It functions as a tetrameric protein, in which the monomers interact through a central oligomerization domain (28, 29). The oligomerization domain is flanked by large intrinsically disordered regions (IDRs) that give HMPV P the ability to interact with a variety of binding partners (28). For instance, the C-terminus of the P protein interacts with RNA-bound N protein to chaperone attachment to the polymerase. Additionally, HMPV P maintains a monomeric pool of RNA-free N protein (N^0^) through an interaction involving the HMPV P N-terminus with the C-terminal domain of the N protein (30). The monomeric N^0^ protein can then be used for ribonucleoprotein assembly at sites of replication where the polymerase synthesizes nascent RNA (30). HMPV P also recruits the antitermination factor M2-1 to the polymerase during transcription to bind nascent viral mRNA (31). Structural analysis of the HMPV polymerase/P protein complex showed the versatility of P monomer interactions with the polymerase, suggesting that P protein IDRs modulate a variety of polymerase functions as well (32). Beyond transcription and replication, HMPV P has been shown to play a role in direct cell-to-cell spread of infection by interacting with actin, or an actin-binding protein, to reorganize the host cell cytoskeleton (33).

This is the first report to analyze phase separation for HMPV IBs. We utilize cellular and purified protein systems to analyze LLPS of HMPV proteins to support the characterization of HMPV IBs as liquid organelles and to determine the interactions required for phase separation. We report that HMPV IBs are liquid-like membrane-less structures that rely on N/P protein interactions. Our *in vitro* data shows that HMPV N and P undergo phase separation and colocalize within liquid droplets when they are mixed in solution. In contrast to MeV and RSV, the HMPV P protein undergoes phase separation in the absence of other viral protein binding partners *in vitro*, suggesting that the P protein may be the key protein to mediate protein interactions to promote IB formation during infection. WT RNA-bound N protein rings formed aggregates in solution but incorporated into liquid droplets in the presence of P protein. These findings suggest for the first time that HMPV P acts as a scaffold protein to support multivalent interactions with HMPV N to promote phase separation and IB formation.

## MATERIALS AND METHODS

### Construction of a recombinant HMPV-mCherryP virus

The plasmids encoding the full-length genome sequence of HMPV strain JPS02-76 (p+JPS07E2) and the accessory proteins N, M2-1, L and P (pCITE-76N, -76M2-1, -76L and -76P) were kindly provided by Dr. Makoto Takeda (National Institute of Infectious Diseases, Tokyo) (34). To insert the mCherryP cassette within the p+JPS07E2 plasmid, a vector containing the partial sequence of N followed by the N-terminus mCherry tagged P sequence, flanked by NheI and SacI restriction sites, was synthesized (GenScript). The sequence within this vector was then subcloned into p+JPS07E2 using the NheI and SacI restriction sites. The correct insertion of the cassette into the plasmid was verified by sequencing. To rescue the recombinant HMPV virus, the methodology described by Shirogane et al. (34) was used. Briefly, BSR cells stably expressing the T7 RNA polymerase were transfected with plasmids p+JPS07E2, pCITE-76N, -76M2-1, -76L and -76P using lipofectamine3000, following manufacturer instructions. Forty-eight hours post-transfection the cells were scrapped from the plate onto the media, and half of the volume overlayed onto a monolayer of Vero cells, in Optimem with TPCK-Try 0.3 µg/mL. Media was replaced every other day until extensive cytopathic effect and fluorescent signal was observed. Cells and media were then recovered and used to propagate the passage 1 of the recombinant virus in Vero cells, as previously described.

### Fluorescence recovery after photobleaching (FRAP)

Vero cells were transfected with pCAGGS plasmid expressing mcherry-P only (P) or co-transfected with pCAGGS plasmids encoding mCherry-P and N protein (P+N). Twenty-four hours post transfection, live cell confocal microscopy was used to perform FRAP at 37°C on punctate regions by drawing a region of interest (ROI) representing a whole inclusion body or an equivalent area in the cytosol with P protein only. Imaging was completed on the Nikon A1R confocal, using a Plan Fluor 40x Oil DIC objective. For photobleaching a laser wavelength of 405nm with a laser power setting of 100% was utilized. Each experiment used 5 seconds of pre-bleaching acquisition, with 4-5 minutes of recovery.

### Expression and purification of HMPV P

The CAN97-83 HMPV P construct was cloned into the plasmid pET 302/NT-His between the cleavage sites EcoRI and XhoI and expressed in BL21(DE3) CodonPlus RIL cells (Agilent) overnight at 37 °C in terrific broth (TB) containing ampicillin after induction at an optical density (OD) of 1.4 with 1 mM isopropyl-β-d-thiogalactopyranoside (IPTG). Cells were lysed with 20 mM Tris, 200 mM NaCl, pH 7.5 containing cOmplete EDTA-free protease inhibitor cocktail (Sigma) and 125 µg/mL lysozyme. After incubating on ice for 20 min, the solution was sonicated three times at 60% intensity for 15 sec. The lysate was spun at 18,000 rpm for 30 min at 4 °C. The crude lysate rocked with HIS-select nickel affinity gel resin (Sigma) for 45 min at 4 °C. The resin was washed one time with lysis buffer and two times with 20 mM Tris, 200 mM NaCl, 20 mM imidazole, pH 7.5. The protein was eluted with 20 mM Tris, 200 mM NaCl, 250 mM imidazole, pH 7.5. The eluate was loaded onto a HiTrap Q HP anion exchange chromatography column (Cytiva). The column was washed with 20 mM Tris, pH 7.5. Then, fractions were eluted with 20 mM Tris, 1 M NaCl, pH 7.5. The fractions containing HMPV P were concentrated and buffer exchanged into 25 mM HEPES, 150 mM KCl, pH 7.5 using a PD-10 desalting column with Sephadex G-25 resin (GE Healthcare).

To reduce nucleic acid binding, some HMPV P lysates were treated with Benzonase during the cell lysis step. Instead of anion exchange, the HIS-select purification was followed by heparin purification using a HiTrap Heparin HP column with an increasing NaCl gradient from 200 mM to 1M prior to buffer exchange with the PD-10 column. After buffer exchange, the protein was concentrated, flash frozen, and stored at −80°C.

### Expression and purification of HMPV N^0^-P

The CAN97-83 HMPV N^0^-P construct with a 6X C-terminal His6-tag was synthesized by GenScript in the pET-29b(+) plasmid and cloned between the NdeI and KpnI cleavage sites. The construct was expressed in *E. coli* Rosetta 2(DE3) competent cells (Novagen) overnight at 18°C in TB containing kanamycin after induction at OD 0.8 with IPTG. Cells were lysed (20 mM Tris, 500 mM NaCl, 10 mM imidazole, pH 7, protease inhibitor, lysozyme, 250 units of Benzonase (Sigma)) and incubated on ice for 20 min. The solution was sonicated and spun as described above except the lysate spun for 45 min. The crude lysate was incubated with resin as described above. The resin was washed once with 20 mM Tris, 500 mM NaCl, 10 mM imidazole, pH 7, twice with 20 mM Tris, 500 mM NaCl, pH 7, and the protein was eluted with 20 mM Tris, 500 mM NaCl, 300 mM imidazole, pH 7. The eluate was concentrated and buffer exchanged into 25 mM HEPES, 150 mM KCl, pH The protein was concentrated and stored as described above.

### Expression and purification of HMPV N-RNA

The CAN97-83 HMPV N construct with a 6X C-terminal His6-tag was synthesized and cloned into pET-29b(+) as described above. The construct was expressed and induced as described for N^0^-P. Cells were lysed (25 mM Tris, 1 M NaCl, pH 8, protease inhibitor, lysozyme, Benzonase) and treated as described above except the lysate spun for 1 hr. The crude lysate was loaded onto a column containing pre-equilibrated resin at 4°C, washed two times (25 mM Tris, 1 M NaCl, pH 8), and eluted (25 mM Tris, 1 M NaCl, 400 mM imidazole, pH 8). The eluate was concentrated and the NaCl concentration of the sample was adjusted to 100 mM using 25 mM Tris, pH 8. Then, the sample was loaded onto a HiTrap Heparin HP column (Sigma) using an increasing NaCl gradient from 200 mM to 1M. Fractions containing the HMPV N protein were buffer exchanged and stored as described above.

### Expression and purification of HMPV N K171A/R186A

The CAN97-83 HMPV N mutant was generated using QuikChange site-directed mutagenesis in pUC57 and subcloned into pET29b(+) using BamHI and XbaI cleavage sites. The construct was expressed, induced, lysed, and spun as described for N^0^-P. The crude lysate was loaded onto a column containing resin as described for N-RNA, and the resin was washed once with 20 mM Tris, 500 mM NaCl, 10 mM imidazole, pH 7 and once with 20 mM Tris, 500 mM NaCl, pH 7. The protein was eluted with 20 mM Tris, 500 mM NaCl, 300 mM imidazole, pH 7. The eluate was concentrated and the NaCl concentration of the sample was adjusted to 100 mM using 20 mM Tris, pH 7. Then, the sample was heparin purified as described for N-RNA. Fractions were buffer exchanged and stored as described above.

### Protein labeling

Prior to buffer exchange, purified HMPV N^0^-P was labeled with Alexa 488 TFP ester (ThermoFisher). The Alexa 488 TFP ester was prepared with DMSO to make a 10 mg/mL solution. The solution was added dropwise to the protein sample. The sample rocked for 1 hr in the dark and was buffer exchanged and stored as described above. Anion exchange purified HMPV P was labeled in a similar manner using Alexa 594 NHS ester (ThermoFisher).

### Droplet assay

A 20% dextran solution was prepared in 25 mM HEPES, 150 mM KCl, pH 7.5. DTT was added to the dextran solution to give a final concentration of 1 mM. HMPV protein constructs were diluted in the 20% dextran, 1 mM DTT, 25 mM HEPES, 150 mM KCl, pH 7.5 solution in 1.5 mL Eppendorf tubes. This solution was used in samples for standard droplet imaging, fusion droplet imaging, and in turbidity assays. For the HMPV P samples tested at different KCl concentrations, similar buffers were prepared with KCl ranging from 0 mM to 500 mM. 1.5 µL of sample was plated on an 8-well printed microscopy slide and covered with a glass coverslip. For droplets imaged at later time points, the slides were stored in a humidified chamber.

### Droplet microscopy imaging

HMPV purified protein samples were imaged using either DIC or epifluorescence on a Nikon Eclipse E600 with the 60X objective. Fusion time lapse images were acquired with MetaMorph software using DIC on a Zeiss Axiovert 200M with the 100X oil objective. Images were acquired at 0.3 sec or 0.5 sec intervals.

### RNA oligomer

The fluorescent RNA decamer was purchased from Integrated DNA Technologies. It was terminated with OH at the 5’ end and 6-carboxyfluorescein at the 3’ end.

### Turbidity assay

Protein solutions were mixed with 20% dextran, 1 mM DTT, 25 mM HEPES, 150 mM KCl, pH 7.5 in clear 96-well plates. The final concentration of the protein was 40 µM. The absorbance of the solutions was measured on a SpectraMax iD3 at 395 nm (21). Readings were taken at 5 min intervals for 8 hr or longer.

### Live cell imaging

VeroE6 cells were seeded in 12 well glass-bottom culture plates and the day after infected with HMPV-mCherryP using a MOI of 3. Cells were kept at 37°C in a 5% CO_2_ atmosphere until imaging. Images were acquired in a LionHeartFX fluorescence microscope using a 60X oil immersion objective. At 24, 48 and 72 hpi infected cultures were imaged for 10 minutes, with images taken every 30 seconds. At least 5 different infected cells were imaged per condition. Alternatively, Vero cells were electroporated with 100 ng of a plasmid encoding mCherryP using a Neon system (ThermoFisher), pulsed at 220V and 950 µF, and subsequently seeded in 6 well glass bottom culture plates. Twenty-four hours post electroporation, cells were infected with rgHMPV at a MOI of 3 and kept for another 48 hours at 37°C in a 5% CO_2_ atmosphere. Cells were imaged in a NikonA1 confocal microscope, acquiring images every 25 seconds using the 60X oil immersion objective.

## RESULTS

### HMPV P localizes to liquid-like IBs in transfected and infected cells

During HMPV infection, incoming and newly synthesized ribonucleoproteins concentrate together in the cytoplasm in an actin-dependent manner (10, 35). Eventually, the coalescence of these structures induces the formation of IBs where viral RNA, viral mRNA, P protein, and N protein are detected (10). Inhibition of actin polymerization significantly reduces HMPV genome transcription and replication, suggesting that IB coalescence enhances the efficiency of these processes (10). To gain insights into IB dynamics in HMPV-infected cells, we generated a recombinant virus with a N-terminus mCherry tagged P protein. mCherry-P retained at least 60% of activity in minireplicon assays, while tagging P on its C-terminus resulted in deleterious effects (data not shown). To characterize the growth kinetics of the recombinant HMPV-mCherryP virus, Vero cells were infected at a MOI of 0.1 PFU/cell. Infected cells were maintained in presence (TPCK+) or absence (TPCK-) of trypsin until 16 days post-infection. Viral titers from supernatants increased until day 10, after which virus growth reached a plateau (FIG. 1A). As expected, HMPV did not grow efficiently in the absence of TPCK trypsin (FIG. 1A). Viral titers for the recombinant HMPV-mCherryP virus were slightly lower than what previously reported for the recombinant JPS02-76EGFP virus (Shirogane, 2008), but this was expected since the mCherryP protein did not retain full replicative activity. It was previously shown that IBs coalesce to a small number of larger structures over the early part of infection, and that this process correlates with maximization of replication efficiency (10). In agreement with this, low numbers of IBs were detected in HMPV-mCherryP infected cells at 24 – 72 hours post infection (hpi) (FIG. 1B). In addition, the size of IBs was shown to nearly double from 2 µm to almost 4 µm from 24 to 72 hpi (FIG. 1C), suggesting a maturation of IBs and potentially increased replication during this period. Using live cell imaging, frequent events of fusion and fission between IBs were observed (FIG. 1D, 1E, 1F). The frequency of fission events per cell significantly increased from 24 to 72 hpi (FIG. 1E), coinciding with the increase in size observed at these hpi. Additionally, incorporation of mCherryP into IBs was observed in mCherryP electroporated - rgHMPV infected cells (FIG. 1G). Fusion and fission events of the IBs in these transfected-infected cells was observed using live-cell imaging, suggesting that HMPV P has an inherent propensity to be incorporated into IBs. Altogether, our results suggest that as infection progress, HMPV IBs grow and increase in complexity and dynamic behavior.

**Figure 1.**
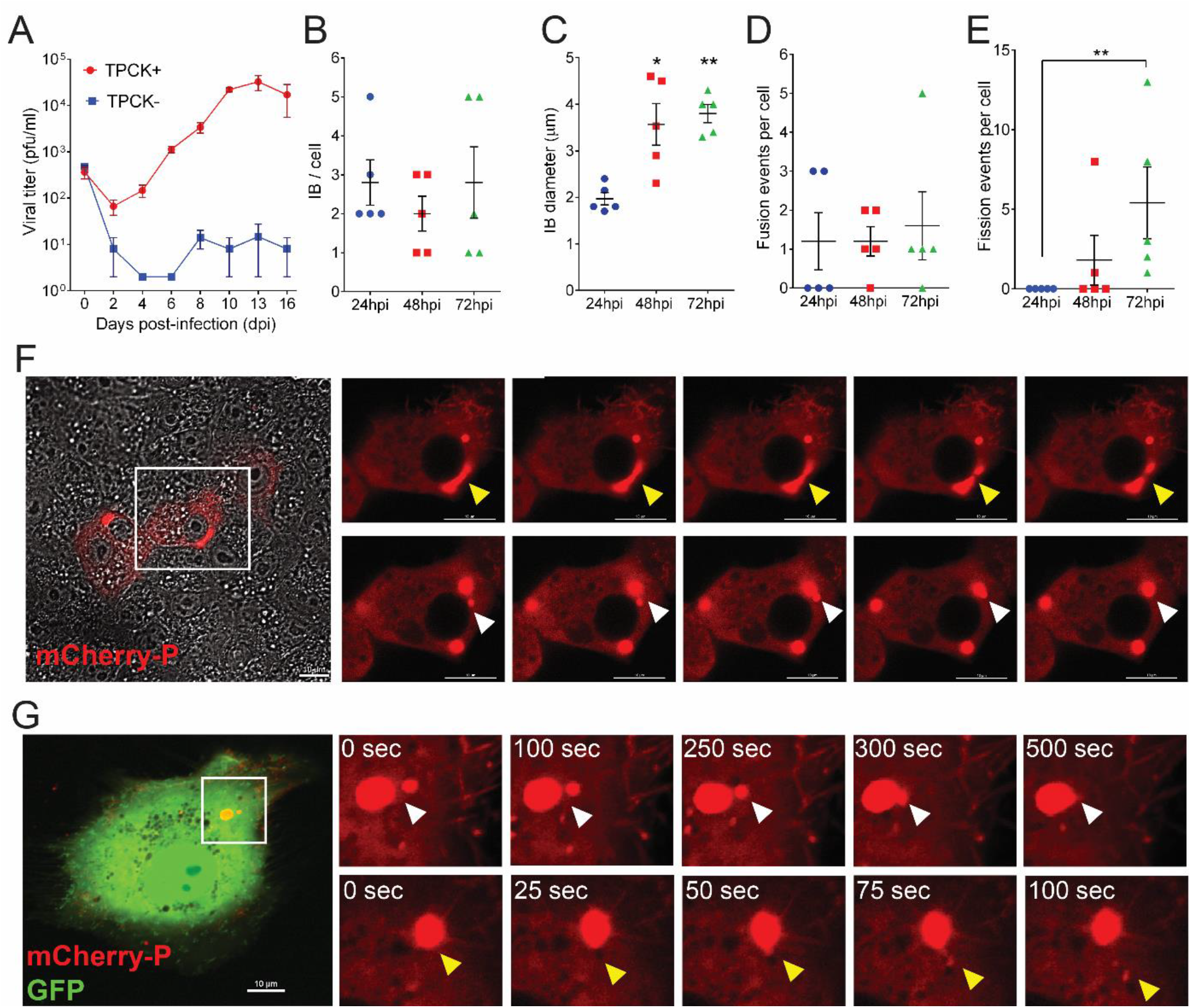
Characterization of a recombinant HMPV-mCherryP virus. (A) Vero cells were infected with a MOI of 0.1 and cells were kept in absence (TPCK-) or presence (TPCK+) of trypsin until day 16 post-infection. Virus was harvested from the cell supernatants every other day and titrated. Vero cells were infected with HMPV-mCherryP virus using a MOI of 3 to quantify the number of IBs per cell (B) and IB diameter (C) at different times post-infection. (D) Fusion and (E) fission events were counted in Vero cells infected with HMPV-mCherryP at a MOI of 3, during a lapse of 10 min. Images were acquired every 30 sec using a LionHeartFX fluorescence microscope. (F) Time-lapse microscopy of Vero cells infected with HMPV-mCherryP, highlighting fission events (upper panels, yellow arrowheads) and fusion events (lower panels, white arrowheads). (G) Vero cells were electroporated with a plasmid encoding mCherryP and subsequently infected with rgHMPV-GFP virus. Fourty eight hours post infection time-lapse microscopy was performed using a NikonA1 confocal microscope, with images acquired every 25 sec. Fusion (white arrowheads) and fission (yellow arrowheads) events are shown. Statistical analysis was performed using Student’s t-test. *P<0.1; **p<0.01.

The liquid-like nature of HMPV IB-like structures was analyzed in transfected cells using FRAP to compare fluorescence recovery rates. When cells were transfected with HMPV P alone, the P protein showed diffuse cytosolic localization and rapid fluorescence recovery (FIG 2). Alternatively, when cells were transfected with both HMPV N and P to induce IB-like structure formation, HMPV P fluorescence recovery rates in the region of the IB were reduced but recovery was still observed, consistent with what is expected for membrane-less liquid-like organelles. This suggests that interactions between HMPV N and P lead to changes in cellular protein dynamics to form phase separated regions. Together, these results support the characterization of HMPV IBs as liquid organelles formed by LLPS as sites for efficient replication and transcription.

**Figure 2.**
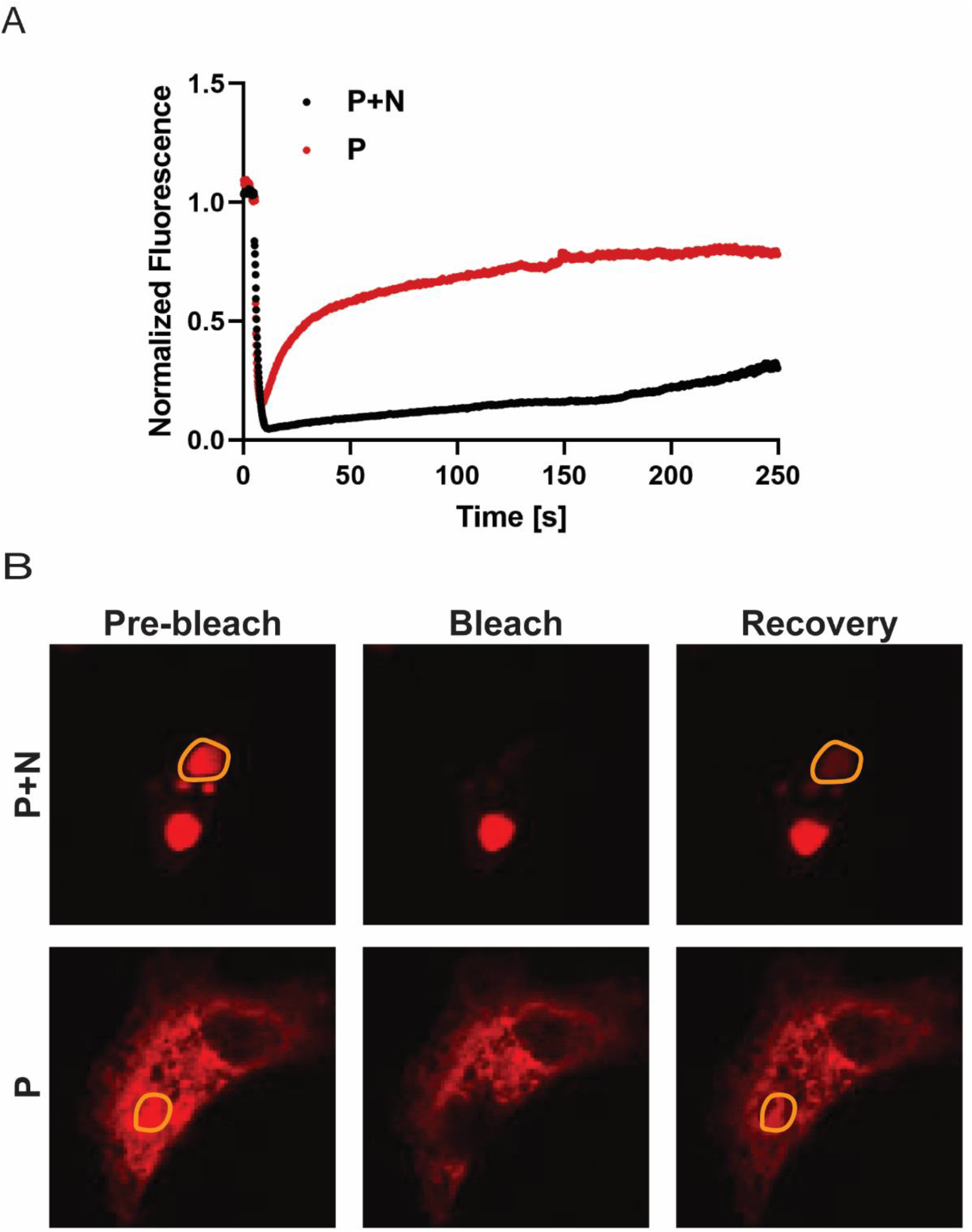
FRAP analysis of HMPV P protein in inclusion bodies and the cytosol. (A) Vero cells were transfected with pCAGGS plasmid expressing mcherry-P only (P) or co-transfected with pCAGGS plasmids encoding mCherry-P and N protein (P+N). 24 hours post transfection, live cell confocal microscopy was used to perform FRAP at 37°C on punctate regions by drawing a region of interest (ROI) representing a whole inclusion body or an equivalent area in the cytosol with P protein only. FRAP data were corrected for background, normalized and are represented as means from the recovery curves. (B) Live cell confocal images collected during FRAP, showing recovery profiles of inclusions 4 min post-bleaching. Bleaching was performed at 100% laser power.

### HMPV P phase separates independently *in vitro*

Since the HMPV N and P proteins are the minimum requirements for IB-like structure formation in eukaryotic cells, recombinant versions of the proteins were expressed in *E. coli* and purified for *in vitro* analysis. Full-length, His_6_-tagged HMPV P was purified by immobilized metal affinity chromatography (IMAC) followed by anion exchange chromatography. Purified HMPV P was then tested in the presence of the crowding agent dextran to assess its ability to undergo LLPS. LLPS is typically driven by scaffold proteins with specific features that promote multivalent interactions with other proteins or RNA (36–38). HMPV P, which includes long IDRs and alternating charged regions, fits the criteria of an LLPS scaffold protein (28). Unlike the reports for MeV and RSV, purified HMPV P formed liquid droplets in the absence of N that were visualized using differential interference contrast (DIC) microscopy, and droplet formation was dependent on the concentration of the P protein (FIG. 3A). Time lapse imaging of the HMPV P droplets showed that they underwent fusion, consistent with the idea that they possess a liquid nature (FIG. 3B). A turbidity assay was also used to analyze purified HMPV P phase separation. The absorbance of the purified HMPV P protein solution was measured at 395 nm at different time points to detect LLPS. The measurements showed a peak for the absorbance above 0.12 between two and four hours, supporting the microscopy imaging results that HMPV P phase separates in the absence of other viral protein binding partners (FIG. 3C).

**Figure 3.**
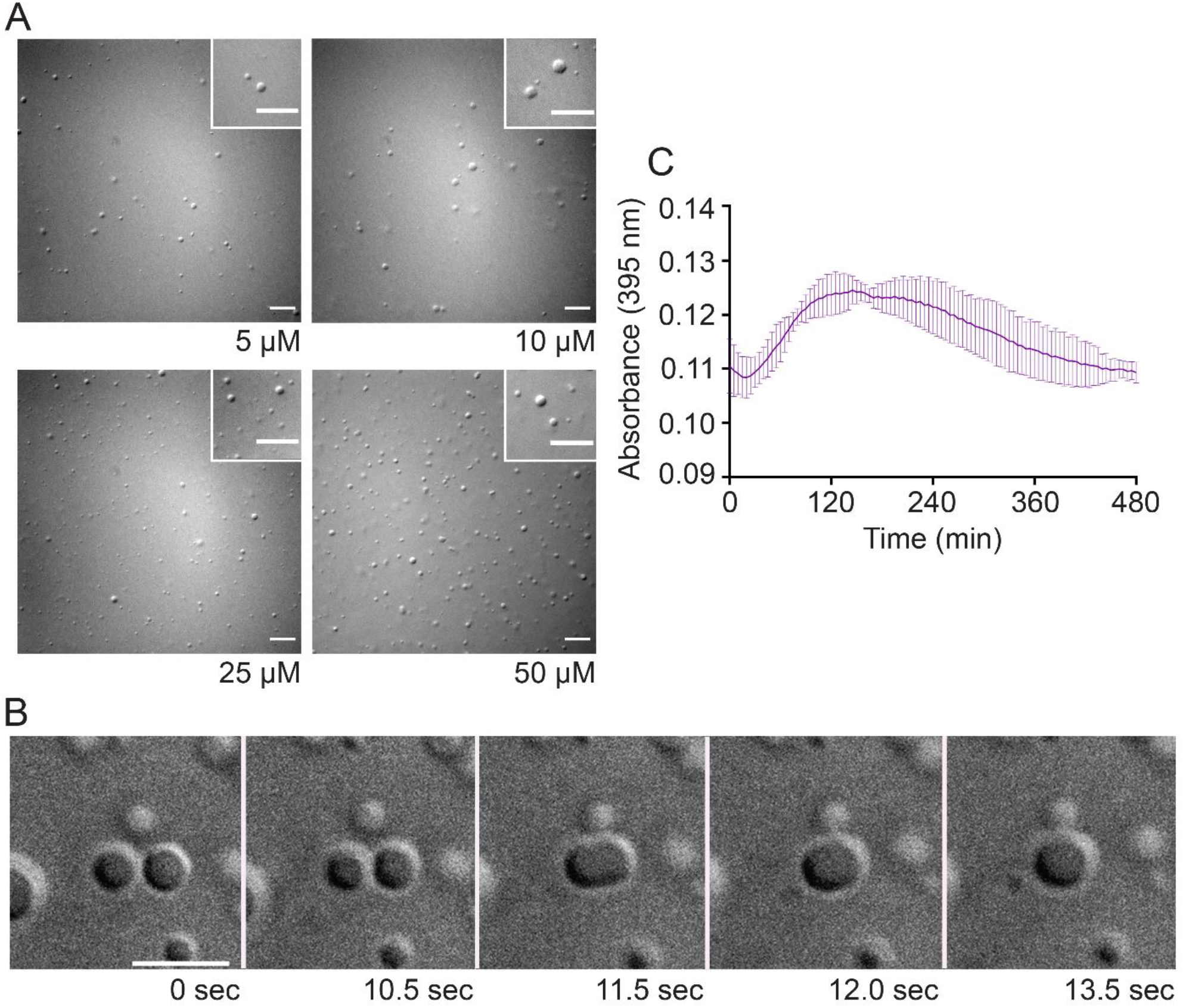
Anion exchange purified HMPV P phase separates independently *in vitro*. (A) Anion exchanged purified HMPV P was tested at concentrations ranging from 5 µM to 50 µM in a droplet assay (maximum droplet size = 3.4 µm). DIC microscopy imaging of droplets was performed with a 60X objective on a Nikon Eclipse E600. The scale bar is 10 µm. (B) Time lapse imaging of anion exchange purified HMPV P (80 µM) droplet fusion was acquired using a 100X oil objective on a Zeiss Axiovert 200M microscope. The scale bar is 5 µm. (C) Anion exchange purified HMPV P (40 µM) was mixed with turbidity assay buffer in a clear 96-well plate. Thesolution was analyzed using a SpectraMax iD3 to measure the absorbance at 395nm at 5 min intervals with mixing.

### Interactions with nucleic acid modulate HMPV P phase separation dynamics

Using the protein purification protocol described above, we noticed that the A260/280 ratio was approximately 1.08, suggesting that the HMPV P protein sample contained nucleic acid. Since nucleic acids are known to play a role in LLPS, we utilized an alternative purification protocol to determine if removing the nucleic acid would influence HMPV P liquid droplet formation. The alternative protocol included treatment with Benzonase nuclease and an IMAC purification step followed by a heparin affinity column purification. This method was successful in removing some of the nucleic acid as indicated by the decreased A260/280 ratio of 0.85. Interestingly, DIC microscopy analysis showed that the recombinant HMPV P protein purified by our alternative protocol formed larger liquid droplets than the original protein sample (FIG. 4A). In addition, time lapse imaging analysis showed that the liquid droplets were capable of fusing (FIG. 4B). Turbidity assay results for the heparin purified HMPV P protein were similar to previous samples, with a peak above 0.12 between two and four hours (FIG. 4C). These results suggest that the presence of increased levels of nucleic acid modulate HMPV P phase separation dynamics. Charge interactions are known to influence phase separation and nucleic acid binding, so both versions of purified HMPV P (anion exchange purified and heparin purified) were analyzed for liquid droplet formation using buffers with different concentrations of potassium chloride (KCl) ranging from 0 to 500 mM. For the anion exchange purified HMPV P, liquid droplets were easily detected between 150 and 250 mM KCl. However, droplet formation was inhibited at concentrations below or above that range (FIG. 5). In contrast, the heparin purified P protein was able to form droplets with as little as 100mM KCl, and as much as 500mM KCl (FIG 5.). For the heparin purified HMPV P, the largest droplets formed at 150 mM KCl droplet size ranging from 0.5-5μm, with an average of 0.64 μm. Average IB size was lower, comparatively, in 250mM salt (average droplet size of 2.5 μm), and similar results were observed at the 500mM concentration. The anion exchanged protein also formed the largest droplets at the 150mM concentration with an average size of 2.36 μm ranging from 0.57 μm −6.8 μm. Anion purified P protein also formed smaller droplets with increasing salt concentration above 150 mM, with an average diameter of 1.7 μm at 250mM KCl and 1 μm at 300mM KCl. Droplets were consistently smaller for the anion exchange purified protein in contrast to the heparin purified product at comparable salt concentrations. These results suggest that HMPV P protein samples containing higher levels of nucleic acid are more sensitive to changes in charge, thus leading to the disruption of liquid droplet formation *in vitro*.

**Figure 4.**
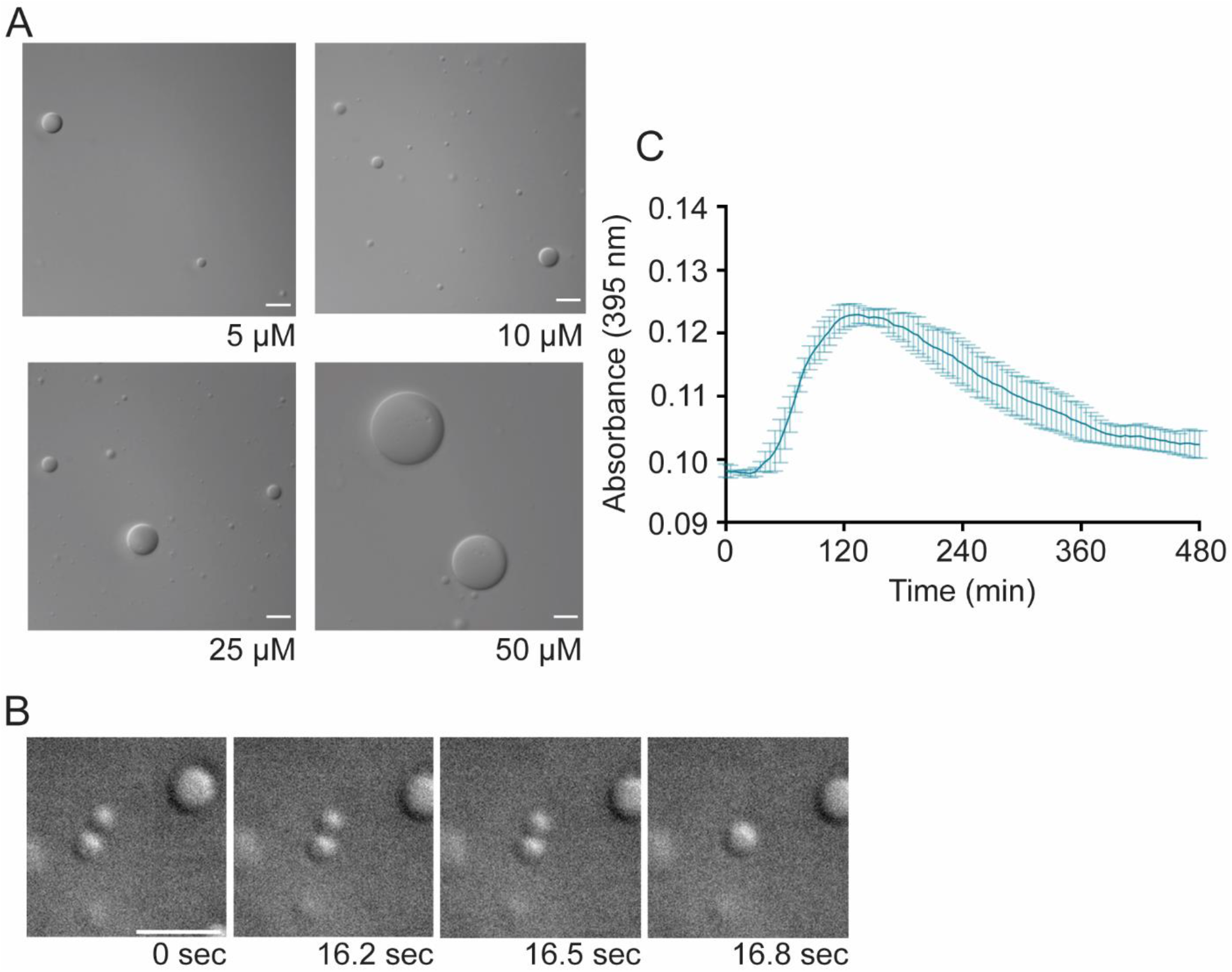
Heparin purified HMPV P phase separates independently *in vitro*. Heparin purified HMPV P was tested at concentrations ranging from 5 µM to 50 µM in a droplet assay (maximum droplet size >50 µm). DIC microscopy imaging of droplets was performed with a 60X objective on a Nikon Eclipse E600. The scale bar is 10 µm. (B) Time lapse imaging of heparin purified HMPV P (150 uM) droplet fusion was acquired using a 100X oil objective on a Zeiss Axiovert 200M microscope. The scale bar is 5 µm. Heparin purified HMPV P (40 µM) was mixed with turbidity assay buffer in a clear 96-well plate. The solution was analyzed using a SpectraMax iD3 to measure the absorbance at 395 nm at 5 min intervals with mixing.

**Figure 5.**
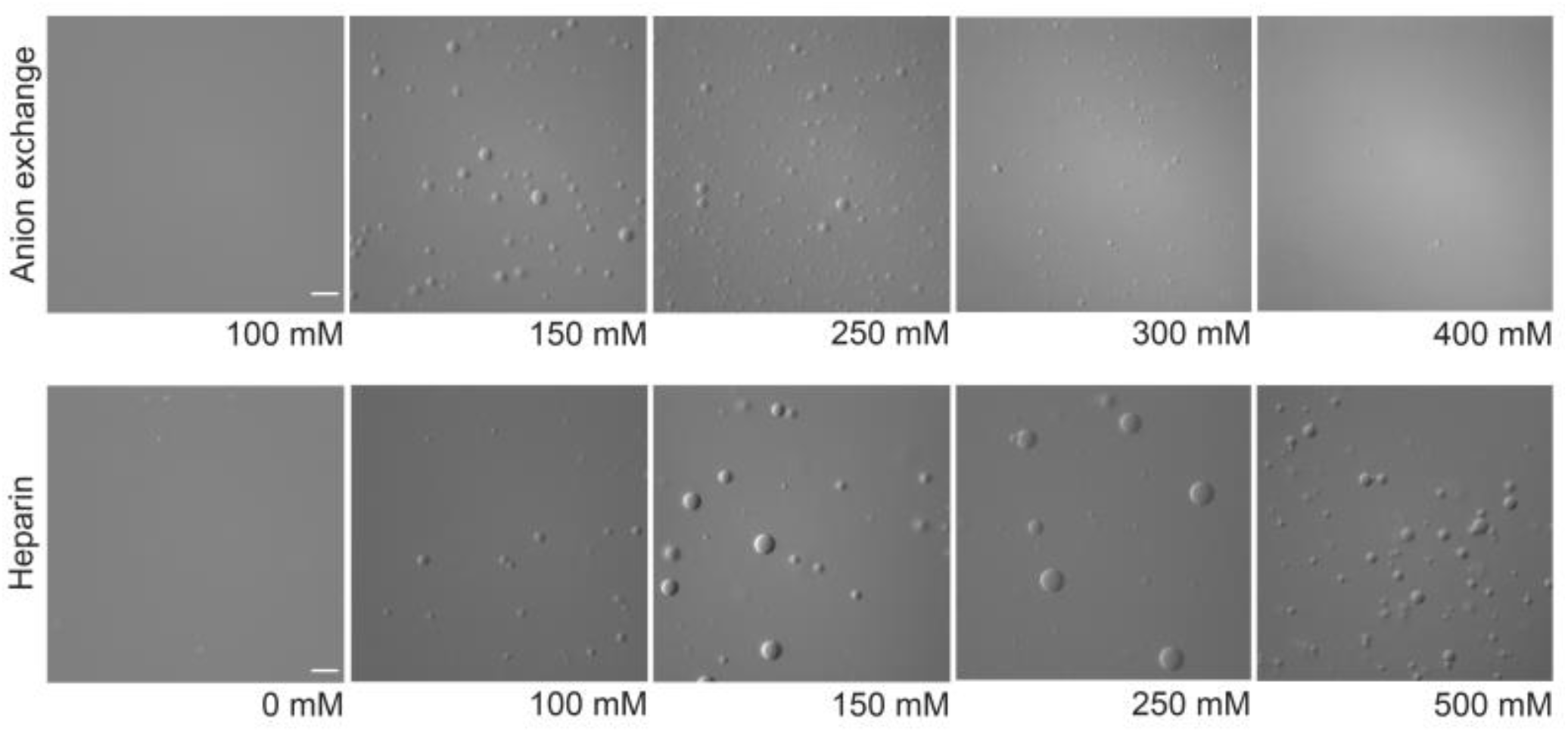
Interactions with nucleic acid modulate HMPV P phase separation dynamics. Anion exchange purified HMPV P (15 µM) and heparin purified HMPV P (15 µM) were tested in a droplet assay using buffers with different concentrations of KCl ranging from 0 mM to 500 mM. DIC microscopy imaging of droplets was performed with a 60X objective on a Nikon Eclipse E600. The scale bar is 10 µm, and the magnification is the same for all images.

### HMPV P recruits N^0^-P to liquid droplets

WT HMPV N spontaneously oligomerizes and binds to nonspecific RNAs during standard purification procedures (30). Thus, we utilized a recombinant N^0^-P construct that includes full-length N (1-394) fused to a P peptide (1–40) to maintain N in a monomeric, RNA-free form for purified protein analysis (FIG. 6A), a strategy that had been successfully utilized by Renner et al. (30). The N^0^-P construct purified by IMAC formed gel-like structures that clumped together in irregular shapes that were visualized by DIC microscopy (FIG. 6B). Unlike anion exchange purified HMPV P, the gel-like HMPV N^0^-P structures remained partially undissolved in 500 mM KCl (data not shown). Over time, these gel-like structures aggregated together but did not undergo fusion (FIG. 6B). In agreement with our microscopy results, turbidity assays performed with the IMAC purified N^0^-P protein gave high absorbance readings that peaked above 0.6, further indicating that the gel-like structures were aggregating in solution (FIG. 6E). The subsequent drop in absorbance suggests that the aggregates settled to the bottom of the 96-well plate.

**Figure 6.**
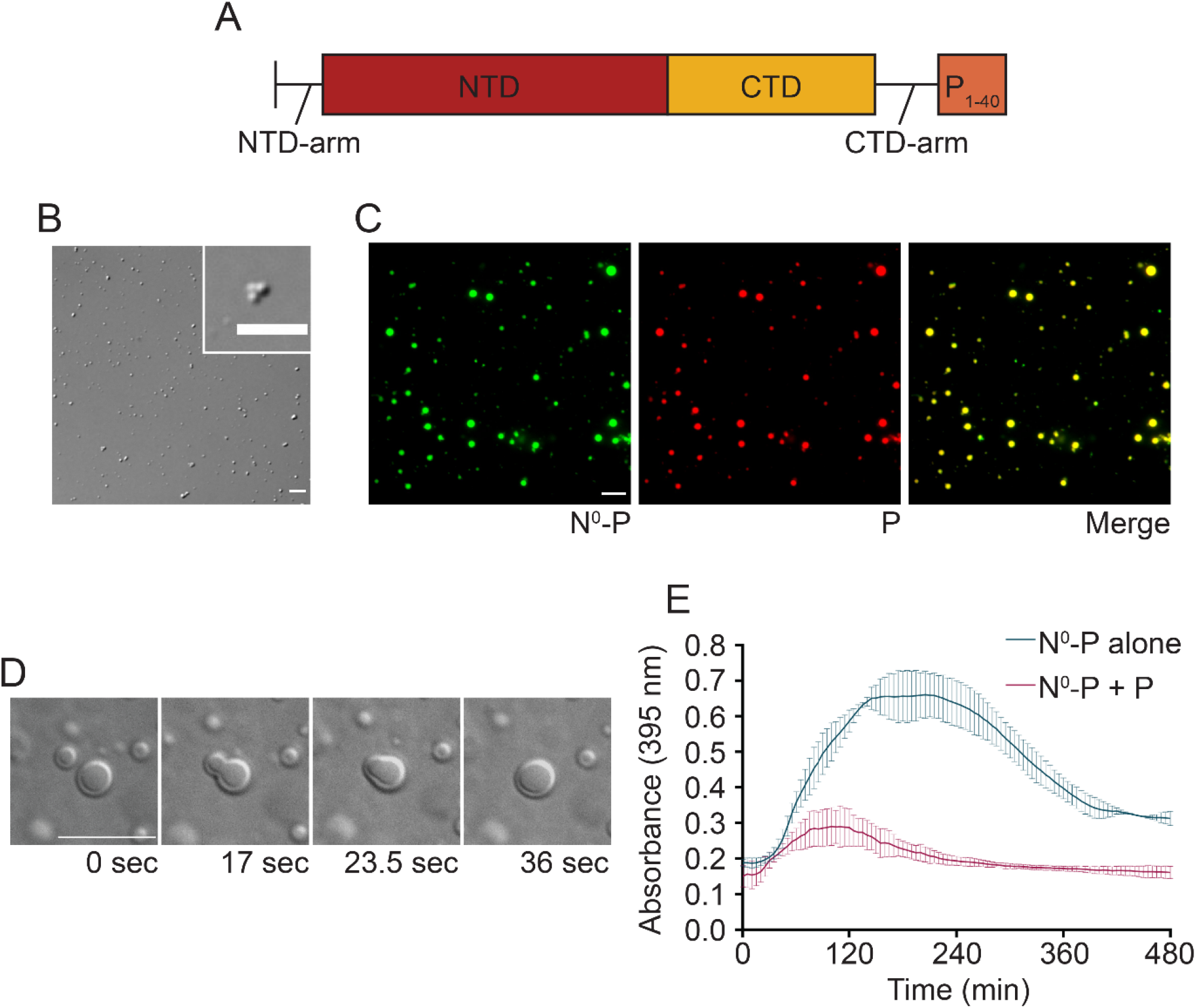
HMPV P recruits N^0^-P to liquid droplets. (A) Schematic of the N^0^-P construct which includes full-length HMPV N fused to the first 40 amino acids of HMPV P. (B) HMPV N^0^-P (15 µM) was tested in a droplet assay. DIC images were acquired at different time points using a 60X objective on a Nikon Eclipse E600. The scale bar is 7 µm. (C) HMPV N^0^-P (15 µM) labeled with Alexa 488 TFP ester was mixed with anion exchange purified HMPV P (15 µM) labeled with Alexa 594 NHS ester in a droplet assay. Fluorescence images were acquired using a 60X objective on a Nikon Eclipse E600. The scale bar is 10 µm. (D) HMPV N^0^-P (50 µM) was mixed with anion exchange purified HMPV P (50 µM). Time lapse imaging of N^0^-P/P droplet fusion was acquired using a 100X oil objective on a Zeiss Axiovert 200M microscope. The scale bar is 10 µm. (E) HMPV N^0^-P (40 µM) was tested alone or with anion exchange purified HMPV P (40 µM) in a turbidity assay. The protein solutions were plated in a clear 96-well plate with turbidity assay buffer, and the absorbance was measured at 395 nm by a SpectraMax iD3 at 5 min intervals.

The N^0^-P construct was examined in combination with anion exchange purified HMPV P using a droplet assay to determine if the P protein could influence N^0^-P dynamics in solution. DIC and fluorescence microscopy analyses showed that mixing the two proteins led to enhanced LLPS as indicated by the presence of larger and more numerous droplets than we previously observed for HMPV P alone. N^0^-P and P were incorporated into the same liquid droplets, as indicated by the colocalization of the fluorescent signals used to label the proteins (FIG. 6C). In addition, the phase separated droplets underwent fusion events (FIG. 6D). A turbidity assay was utilized to determine if combining N^0^-P with P affected the absorbance of the solution. The results showed that compared to N^0^-P alone, the combination of N^0^-P and P led to lower absorbance readings that peaked around 0.3 at two hours (FIG. 6E). Together, these findings support that HMPV P facilitates interactions with N^0^-P to recruit the protein into liquid droplets, and interactions between the proteins prevent the N^0^-P construct from transitioning to a gel-like state.

### HMPV P recruits N-RNA rings to liquid droplets

In addition to the monomeric N^0^-P construct, WT N-RNA rings were purified for LLPS analysis in the presence or absence of HMPV P. DIC imaging of N-RNA rings in the droplet assay showed the formation of clumped, irregularly-shaped structures that did not undergo fusion, suggesting that this protein-RNA complex does not for liquid-like phase separated structures independently (FIG. 7A). Combining the purified N-RNA rings with heparin purified HMPV P resulted in N-RNA complex incorporation into liquid droplets (FIG. 7B). The N-RNA/P droplets (maximum droplet size = 6 µm) were generally smaller than the P alone droplets (maximum droplet size = 11 µm), suggesting that this combination influences phase separation dynamics. The influence of HMPV P on N-RNA for liquid droplet formation was reflected in the turbidity assay results which showed a lower peak for absorbance around 0.2 at two hours, compared to the absorbance for N-RNA alone (FIG. 7D).

**Figure 7.**
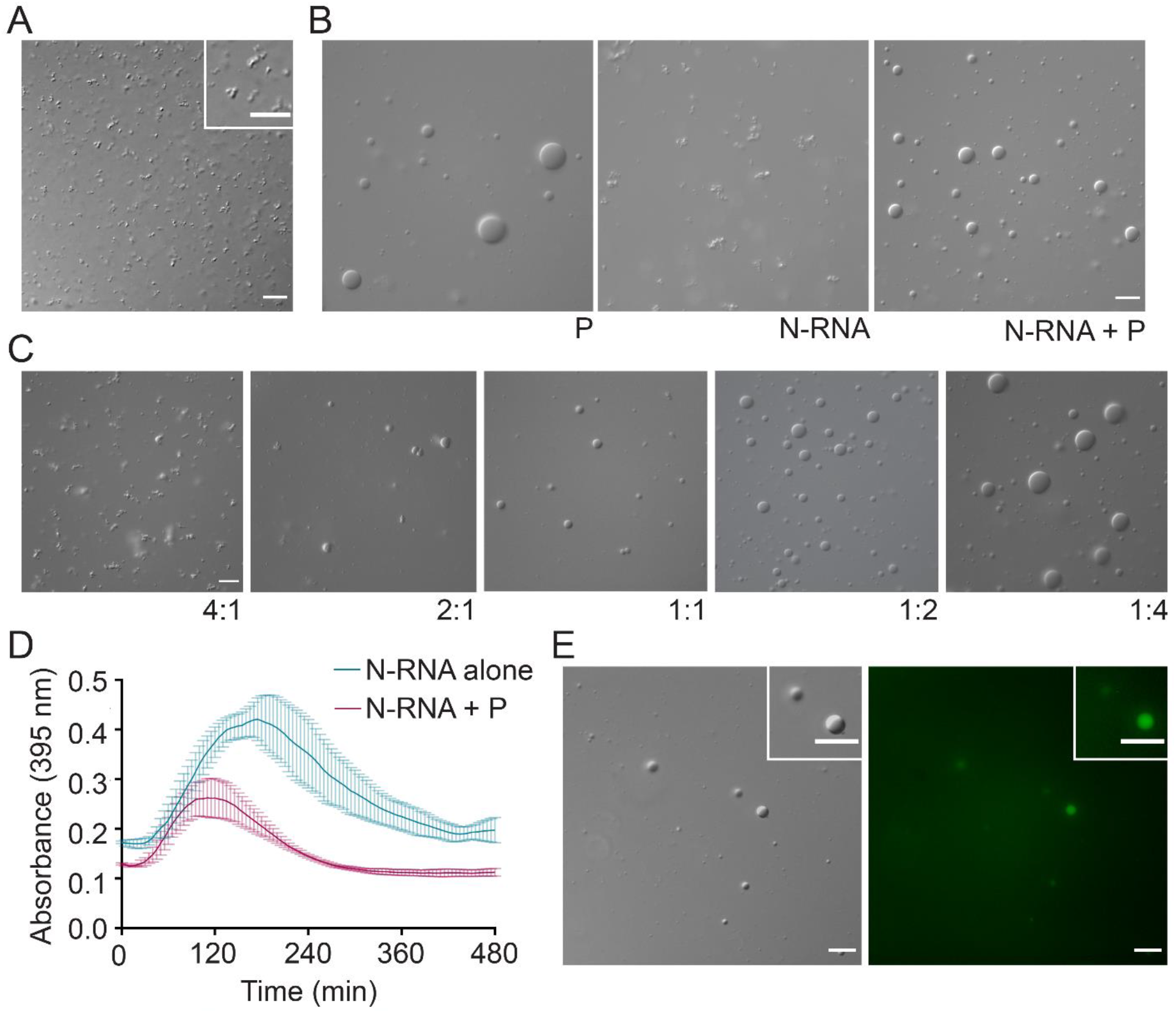
HMPV P recruits N-RNA rings to liquid droplets. (A) HMPV N-RNA (25 µM) was tested in a droplet assay. DIC microscopy imaging of droplets was performed with a 60X objective on a Nikon Eclipse E600. The scale bar is 10 µm. (B) HMPV N-RNA (15 µM) was mixed with heparin purified HMPV P (15 µM) in a droplet assay. DIC images were acquired using a 60X objective on a Nikon EclipseE600. The scale bar is 10 µm. (C) HMPV N-RNA and heparin purified HMPV P were tested in a droplet assay at different ratios (4:1 = 20 µM N-RNA: 5 µM P; 2:1 = 10 µM N-RNA: 5 µM P; 1:1 = 5 µM N-RNA: 5 µM P; 1:2 = 5 µM N-RNA: 10 µM P; 1:4 = 5 µM N-RNA: 20 µM P). DIC microscopy imaging of droplets was performed as described above. (D) HMPV N-RNA (40 µM) was tested alone or with heparin purified HMPV P (40 µM) in a turbidity assay. The protein solutions were plated in a clear 96-well plate with turbidity assay buffer, and the absorbancewas measured at 395 nm by a SpectraMax iD3 at 5 min intervals with mixing. (E) HMPV N-RNA (15 µM), heparin purified HMPV P (15 µM), and an RNA decamer tagged with 6-carboxyfluorescein on the 3’ end (5 µM) were mixed and tested in a droplet assay. DIC and fluorescence microscopy imaging of droplets was performed as described above. The scale bar is 10 µm.

HMPV P and N-RNA were tested in our *in vitro* system at different ratios to determine the conditions that were required for N-RNA to be recruited to liquid droplets. Though N-RNA aggregates were still present at 4:1 and 2:1 ratios of N-RNA/P, round droplets were easily detected at a 1:1 ratio. The number and size of the round droplets increased in samples with a higher proportion of HMPV P (1:2 and 1:4) (FIG. 7C). These results suggest that a specific ratio of N-RNA/P must be met before N-RNA is induced to phase separate into droplets. Liquid droplets containing HMPV P and N-RNA were also shown to incorporate a fluorescent RNA oligomer (FIG. 7E). These findings highlight that HMPV P, N, and RNA form complex multivalent interactions to promote phase separation and to support the structure of IBs required to enhance replication and transcription.

### HMPV P recruits RNA-binding mutant N K171A/R186A to gel-like droplets

An HMPV N mutant (K171A/R186A) was generated to determine if RNA binding affects N recruitment into droplets. This mutant was designed based on the RSV construct N K170A/R185A which lacks the ability to bind RNA (39). HMPV N^0^-P, N-RNA, and N K171A/R186A were tested individually with a fluorescent RNA oligomer in a droplet assay to assess the RNA-binding capabilities of the different constructs. The RNA oligomer incorporated into N^0^-P structures, suggesting that the RNA may disrupt binding of the P_1-40_ peptide to N (FIG. 8A). Alternatively, the P_1-40_ peptide, which contains positively charged residues, may interact directly with the RNA oligomer. In contrast to N^0^-P, N-RNA showed a weak interaction with the fluorescent RNA oligomer, suggesting that oligomer poorly disrupts the existing interactions between HMPV N and RNA in stable ring structures (FIG. 8A). The N K171A/R186A mutant showed no colocalization with the fluorescent RNA oligomer, suggesting that mutation of these residues in the RNA-binding cleft of HMPV N effectively inhibits RNA binding (FIG. 8A).

**Figure 8.**
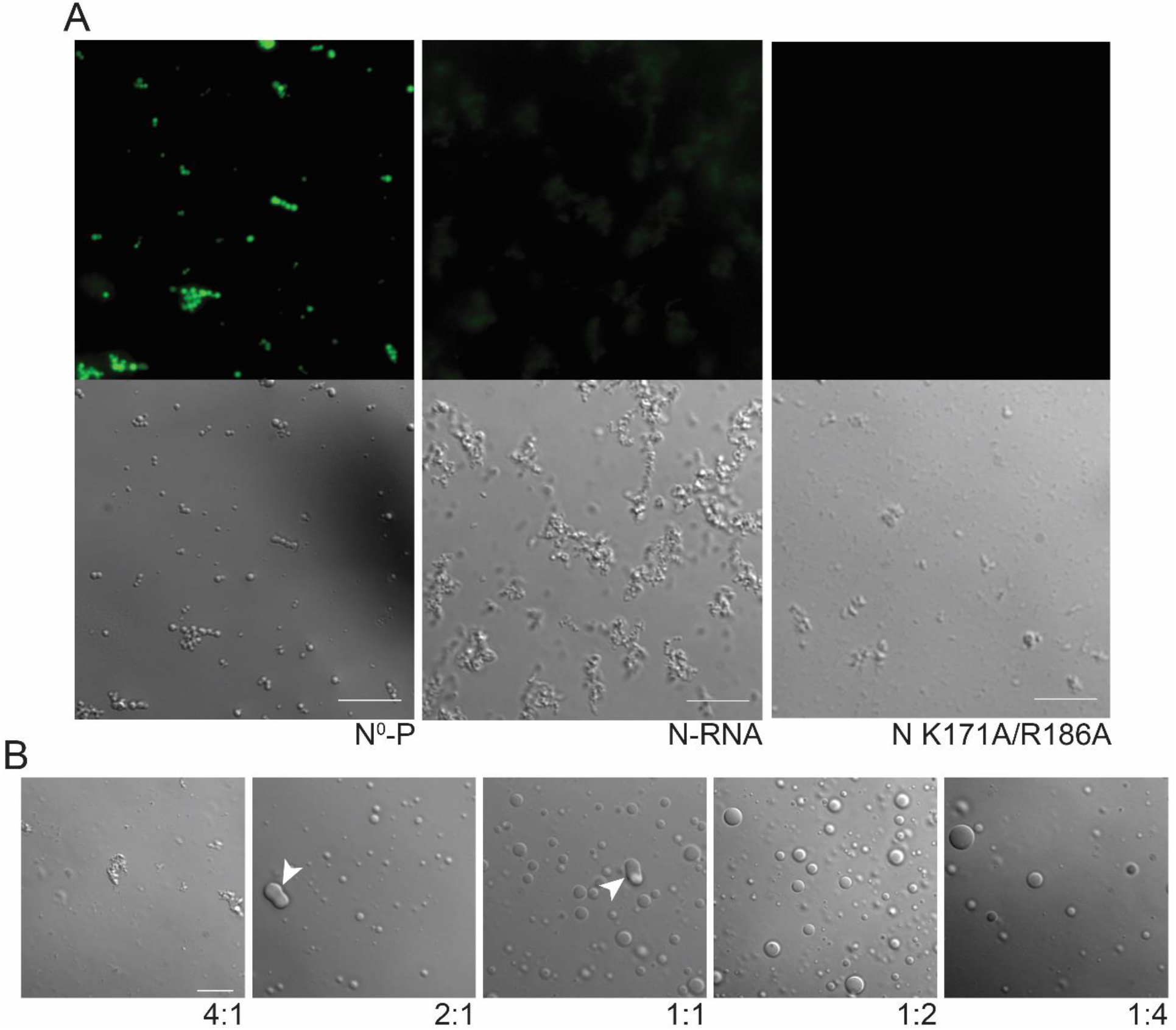
RNA-binding mutant HMPV N K171A/R186A forms gel-like droplets with P. (A) HMPV N^0^-P, N-RNA, or N K171A/R186A (15 µM) were tested in a droplet assay with an RNA decamer tagged with 6-carboxyfluorescein on the 3’ end (5 µM). DIC and fluorescence microscopy imaging was performed on a Zeiss Axiovert 200M with a 63X oil objective. The scale bar is 10 µm. (B) HMPV N K171A/R186A and heparin purified HMPV P were tested in a droplet assay at different ratios (4:1 = 20 µM N K171A/R186A: 5 µM P; 2:1 = 10 µM N K171A/R186A: 5 µM P; 1:1 = 5 µM N K171A/R186A: 5 µM P; 1:2 = 5 µM N K171A/R186A: 10 µM P; 1:4 = 5 µM N K171S/R186A: 20 µM P). DIC microscopy imaging of droplets was performed on a Zeiss Axiovert 200M with a 63X oil objective. White arrowheads indicate altered droplet fusion. The scale bar is 10 µm.

Subsequently, HMPV N K171A/R186A was tested at different ratios with heparin purified HMPV P in a droplet assay to analyze LLPS dynamics (FIG. 8B). At a 4:1 ratio of N K171A/R186A:P, microscopy imaging showed the presence of aggregates and no liquid droplets. Droplets became visible as the concentration of HMPV P was increased relative to the N RNA-binding mutant. However, the droplets exhibited gel-like features, and droplets were seen to adhere to each other (white arrowheads, FIG. 8A) but were unable to undergo fusion. This suggests that RNA binding to HMPV N plays an important role in modulating LLPS dynamics *in vitro* and support our model that HMPV P, N, and RNA form complex interactions in cells to promote replication and transcription.

## DISCUSSION

IB formation has been reported for many negative-strand viruses across the *Mononegavirales* order (40). Recent evidence supports that these structures function as viral factories by concentrating the materials required for replication and transcription. Cellular studies of IBs have shown that these structures are membrane-less and dynamic, which led to the characterization of IBs as liquid organelles. Though membrane-less organelles have been recognized in the cell for decades, scientists have only recently linked the formation of these structures to the process of LLPS. Understanding the role of LLPS in IB formation may be critical for discovering new targets for pan-antiviral development. Until now, no reports have been published to determine if HMPV IBs are consistent with phase separated liquid organelles. Cellular experiments were utilized to analyze the dynamic nature of HMPV IBs in infected or transfected cells. Additionally, *in vitro* experiments were performed to test recombinant versions of HMPV purified proteins in LLPS assays. These studies provide strong evidence for the novel role of HMPV P as a scaffold for recruiting N protein and other components to IBs.

Cellular analysis of HMPV IBs using live cell imaging and FRAP showed that these structures form as distinct phase separated regions that exchange components with the surrounding cytoplasm and undergo fusion and fission. Interestingly, our analysis of cells infected with recombinant HMPV-mCherryP virus showed that levels of IB fusion remained stable from 24 to 72 hpi, whereas fission events significantly increased during this time frame (FIG. 1). Since IB diameter increased significantly from 24 to 72 hpi, this suggests that early fusion events and the incorporation of cellular components, newly synthesized HMPV proteins, and viral RNA contribute to the growth of IBs over time. Furthermore, the plateau of fusion events suggests that IBs likely mature to a gel-like state by 72 hpi. In contrast, fission levels may be less affected by the gel-like state due to the activity of vesicles that traffic IB components out of large IBs, as we have observed via live cell imaging (data not shown). IB fission events may be linked to the release of small replication bodies to create new viral factories (11). Alternatively, fission may promote the formation of unique IB subpopulations later in infection to facilitate assembly and budding (41). These findings highlight the crucial role IBs likely play in establishing and promoting the spread of infection.

To analyze the protein interactions that govern IB formation, we compared purified HMPV P in the presence or absence of nucleic acid with a set of N forms using *in vitro* LLPS experiments. Importantly, we found that HMPV P phase separates independently *in vitro* and can act as a scaffold to recruit other client proteins, including HMPV N, to liquid droplets to drive LLPS. Additionally, our findings suggest a previously undescribed role for HMPV P in interacting with RNA to modulate phase separation. As the genome and antigenome of HMPV are fully coated by N, P could interact with viral mRNAs or with cellular RNAs in the context of an infection. Features of viral P proteins, including long regions of intrinsic disorder, match the molecular signature of proteins that phase separate under physiological conditions (24, 28). Furthermore, they are consistent with HMPV P as a scaffold protein that binds a variety of substrates, such as viral proteins and RNA, to promote IB formation. The recently published structure of the HMPV polymerase/P protein complex highlights the importance of IDRs in allowing HMPV P to adopt a variety of binding conformations (32, 42). The propensity for HMPV P to mediate multivalent interactions and phase separate independently *in vitro* suggests that it regulates similar functions for IB formation during infection (36).

In contrast to our *in vitro* results, the HMPV N and P proteins must be co-expressed in cells to generate IB-like structures (26). Without HMPV N, the P protein showed both diffuse cytoplasmic localization and peripheral filopodia-like localization (26). This difference between the cellular and *in vitro* systems suggests that host factors in the cytoplasm may block HMPV P interactions required to induce LLPS. Co-expression of HMPV P and N likely acts to initiate LLPS in cells by concentrating enough IB components to drive phase separation. LLPS is a highly sensitive process that depends on factors such as protein/RNA concentration, salt content, post-translational modifications, pH, and temperature (24). One or more of these factors may prevent HMPV P from phase separating when expressed independently in cells. Furthermore, results showed that removal of nucleic acid from purified HMPV P modulated liquid droplet formation *in vitro* (FIG. 5), suggesting that HMPV P phase separation in cells may be significantly impacted by the presence of RNA. Additionally, during HMPV infection, N protein is always expressed in excess compared to HMPV P due to the location of the N gene within the viral genome. This suggests that HMPV P may lack opportunities for independent phase separation during infection due the local concentration of other viral factors involved in IB formation. Though *in vitro* studies are crucial for deciphering the mechanisms of HMPV phase separation, the increased complexity of protein and RNA interactions during cellular infection must always be considered.

Our work is the first to provide evidence of a viral P protein functioning as a phase separation scaffold, in contrast to related systems. Recent reports on MeV and RSV showed that a combination of the N and P proteins was required to induce droplet formation *in vitro* (21, 22). Though *Mononegavirales* P proteins share common structural features, they lack sequence similarity and vary in length (43). MeV P is 213 residues longer than HMPV P and includes a folded C-terminal (XD) domain after the unfolded P_loop_. The pneumoviral RSV and HMPV P proteins possess similar domain organization, but sequence differences likely promote unique LLPS interactions for each virus. For instance, HMPV P is 53 residues longer than RSV P and contains insertions in the N-terminal and C-terminal domains that may influence IDR behavior (43). The C-terminal domain and oligomerization domain of RSV P were required for liquid droplet formation with N-RNA, suggesting that the acidic insertion in the HMPV P C-terminal domain may modulate phase separation (22, 43). The differences observed for these viral systems emphasize that LLPS is highly dependent on multivalent interactions mediated by the unique composition of the P protein.

Though HMPV N and P are required for IB formation in cells, the role of different N protein forms in phase separation required further exploration. We compared monomeric N protein (N^0^-P) and N-RNA with N K171A/R186A to analyze the effects of oligomerization and RNA binding on LLPS with the HMPV P scaffold. Though all the N forms were recruited to droplets by HMPV P, the incorporation of N K171A/R186A led to the formation of droplets that failed to undergo complete fusion, suggesting that they were gel-like rather than liquid-like in nature. This highlights that RNA interactions with HMPV N and P alter phase separation dynamics and suggests that viral RNA levels may modulate IB maturation during the course of HMPV infection. A minimal MeV LLPS *in vitro* system showed that RNA diffuses into MeV N/P liquid droplets, triggering the formation of nucleocapsid-like particles (21). Interestingly, coexpression of a MeV N RNA-binding mutant with P did not alter the morphology of IB-like structures in cells compared to coexpression of WT N and P (20). These findings suggest that RNA binding is not required for MeV IB formation, but RNA incorporation likely serves to enhance ribonucleoprotein assembly during infection. In contrast, a monomeric RNA-free RSV N mutant failed to form IB-like structures with RSV P in cells, suggesting that RSV N must oligomerize and/or bind RNA to mediate IB formation (22). Our *in vitro* results with HMPV N^0^-P and N K171A/R186A confirmed that N protein oligomerization and RNA interactions are not required for phase separation with HMPV P.

Here, we showed that HMPV IBs are liquid organelles and that HMPV P acts as a scaffold to recruit different forms of N to liquid droplets. We report that nucleic acid interactions with P and N alter phase separation dynamics, suggesting that viral RNA binding plays a significant role in HMPV IB formation and maturation. Recent work on RSV utilized a condensate-hardening drug to block RSV replication in the lungs of infected mice (44). This exciting evidence suggests that IBs of various negative-strand viruses may serve as druggable targets for inhibiting infection. The work presented here builds on the foundation for understanding the formation of IBs and the mechanisms that regulate LLPS for negative-strand viruses. Deciphering the protein and RNA interactions that influence IB phase separation will be essential for the development of pan-antiviral drugs to target viral factories.

## Acknowledgements

We are grateful to Max Renner, Oxford, for critical input on the construction and purification of the various HMPV N protein forms, and to Amanda Wilburn for technical assistance. Funding for this work was provided to NCM by Fondecyt grant 11180269 and to RED by NIAID grant RO1AI40758.

